# Visual processing of manipulable objects in the ventral stream is modulated by inputs from parietal action systems

**DOI:** 10.1101/2025.06.01.657280

**Authors:** Frank E. Garcea, Emma L. Strawderman, William Burns, Matthew Cotroneo, Steven P. Meyers, Tyler Schmidt, Kevin A. Walter, Webster H. Pilcher, Bradford Z. Mahon

## Abstract

Functional object use requires the integration of visuomotor representations processed in the dorsal visual pathway with representations of surface texture and material composition of objects, processed in the ventral visual pathway. Do regions in the ventral visual pathway project outputs to dorsal visual pathway action systems, or are the outputs of the dorsal visual pathway communicated to the ventral visual pathway to modulate processing? And what are the white matter pathways that mediate structural connectivity in support of functional object use? Here we show that the left inferior parietal cortex, a region within the dorsal visual pathway, exerts a direct effect on neural responses in ventral occipital-temporal cortex during visual processing of manipulable objects. We studied a series of consecutively enrolled participants in the pre-operative phase of their neurosurgical care (N = 109) with lesions principally distributed throughout the left hemisphere. Participants completed an object processing category localizer functional MRI experiment in which they viewed images of manipulable objects, animals, faces, and places. We then used Voxel-based Lesion-Activity Mapping (VLAM), a technique in which functional responses in a region-of-interest are used to predict variance in voxel-wise lesion incidence throughout the brain. In the VLAM analyses performed here, we found that lesions to the left anterior intraparietal sulcus and left supramarginal gyrus, two inferior parietal regions known to support object-directed grasping and manipulation, respectively, cause reduced neural responses for manipulable objects (compared to faces, places and animals) in the fusiform gyrus. Parietal lesions do not affect neural responses during visual processing of places in the same region of the fusiform gyrus, even though places elicit stronger responses in the fusiform gyrus than manipulable objects. Seventy-five of 109 participants took part in a common diffusion MRI protocol, permitting a connectome-wide analyses relating white matter fiber integrity to the strength of functional responses for manipulable objects in the left fusiform gyrus. This analysis demonstrated that the descending portion of the left arcuate fasciculus mediates parietal-to-temporal lobe connectivity for manipulable objects, supporting the integration of action representations with conceptual and perceptual attributes of objects. By combining voxel-based and connectome-wide lesion-symptom mapping methods with functional MRI, we have demonstrated that structural connectivity to dorsal visual pathway areas supporting skilled manual action shape category-specific neural responses for manipulable objects within the ventral visual pathway.

## Introduction

Every action performed upon an object – grasping a cup, opening a door, putting on our shoes – requires coordinated processing across a network of temporal, parietal and frontal regions that support functional object grasping and use.^1–4^ The dorsal visual pathway, which projects to the anterior intraparietal sulcus (aIPS), processes volumetric properties and spatial location of objects with respect to the body.^5–8^ Dorsal computations may be sufficient to support volumetric grasping, but praxis (skilled object manipulation) requires the integration of visuomotor processing with object properties (shape, material properties, surface texture) that are computed by the ventral object processing pathway.^9^ For instance, the grip force required to grasp a slippery glass, or the hand posture to grasp an upside-down glass on a counter, depend on perceptual analysis that is supported by occipital-temporal regions^9–11^ (for discussion, see ^12^). The aIPS and supramarginal gyrus (SMG), of the inferior parietal lobule, support object-directed grasping and complex object manipulation (praxis), respectively.^7,13–18^ A key open question is which white matter pathways allow parietal-based action representations to interact with occipital-temporal object representations: how does what the eye *sees* inform what the hand *does*?

Longitudinal fibers within the ventral stream, including the inferior longitudinal fasciculus and inferior fronto-occipital fasciculus, integrate processing along the hierarchy of visual regions—and support both feedforward and feedback processing.^19–21^ Traditionally, models of visual object recognition have emphasized that visual perceptual analysis is driven by processes internal to the ventral stream.^6,22–25^ The ventral stream is defined computationally as a pathway that supports visual object identification, with its outputs constituting important inputs to parietal and frontal regions that calibrate object-directed actions.^18,26–28^

Without challenging the maxim that the ventral stream supports identification, it is also the case that ‘identification’ is not just about ‘labeling’. In the context of object-directed action, there are some action properties that are relevant to action, and which are processed in the ventral stream. ^12,29,30^ For instance, if the action plan requires grasping a cup to take a drink, calibration of grip force will drive perceptual analysis of the material properties of the cup (e.g., is it Styrofoam or ceramic?) to determine if it might collapse under a certain grip force. We propose and test the hypothesis that parietal-based action representations provide inputs to occipital-temporal perceptual processes for manipulable objects.

Visual processing of manipulable objects drives differential neural responses in specific regions of the Ventral Temporal Cortex (VTC), in particular along the collateral sulcus and the medial fusiform gyrus.^31,32^ Those regions are functionally defined as exhibiting differential responses to manipulable objects compared to a range of visual baselines, including faces, places, body parts, animals, and written words (for broader findings regarding neural specificity for visual categories, see ^31,33–40^). Neuropsychological studies have shown that focal lesions along the collateral sulcus can cause selective deficits for knowledge of object surface texture and material properties^41,42^ (for convergent fMRI findings, see ^43,44^), and for of processing manipulable objects compared to non-manipulable objects.^45^ Prior research has also shown that the collateral sulcus and medial fusiform gyrus exhibit differential functional connectivity to the left aIPS and SMG.^1,46–48^ For instance, in one study, the voxel-wise pattern of preferences for manipulable objects in left VTC could predict the voxel-wise pattern of functional connectivity to the left SMG.^49^

Interestingly, there can be *misalignment* in the VTC between category-preferences, as determined by relative amplitudes of BOLD responses, and other measures of neural specificity. For instance, images of places and highly contextualized objects (i.e., landmarks, large non-manipulable objects) differentially drive responses in the parahippocampal gyrus^50,51^, which is anterior to the VTC area exhibiting preferences for manipulable objects.^49,52^ However, even within the collateral sulcus and medial fusiform area that exhibits stronger responses to manipulable objects than faces or animals, places and large non-manipulable objects typically elicit stronger BOLD responses than manipulable objects.^29,32,52^ Thus, on the one hand, the amplitude of BOLD responses shows one pattern (stronger for places than manipulable objects; ^52^), while patterns of repetition suppression^32^, functional connectivity^49^, and lesion evidence^41^ demonstrate various degrees of specificity for *manipulable* objects.

Our proposal is that neural responses in left VTC represent a complex summation of processes that result from an analysis of the visual inputs, and processes associated with inputs from parietal action systems.^32,39^ We refer to this framework as a dorsal-to-ventral model. Here seek causal evidence for the dorsal-to-ventral model, by testing ***i)*** if lesions to parietal regions disrupt functional responses in left VTC for manipulable objects, ***ii)*** and by identifying which white matter pathways mediate the effect of parietal lobe lesions on distal functional response in VTC. We studied a group of 109 individuals with acquired brain lesions, and 55 neurotypical controls, who each completed an fMRI task involving manipulable objects, animals, faces, and places, as well as a diffusion-weighted MRI. We first show that lesion presence in the left SMG and aIPS is associated with reduced neural preferences for manipulable objects in left VTC, extending a prior study in a smaller group of individuals with brain lesions.^29^ We then use connectometry and deterministic tractography to describe the white matter pathways that mediate structural connectivity between the identified parietal regions with left VTC manipulable object processing regions. Candidate fiber pathways that integrate object representations in left VTC with skilled action production systems in the SMG and aIPS include the left arcuate fasciculus^53^, the left parietal aslant tract^54^, the left vertical occipital fasciculus^55,56^, the left middle longitudinal fasciculus^57^, and the left inferior fronto-occipital fasciculus.^58^ Here, we find the descending portion of the left arcuate fasciculus is the key pathway that integrates functional grasping and manipulation processes in the left inferior parietal lobule with object processing in the left VTC.

## Methods

### Participants

One-hundred and fourteen individuals with an acquired brain lesion participated in the study. Participants had normal or corrected-to normal vision and were in the pre-operative phase of their neurosurgical care (49 females; mean age, 46.54 y; SD 16.48; see Supplemental Table 1 for demographic variables and Supplemental Figure 1 for lesion overlap among participants). Participants took part in a T1 anatomical scan, an object processing category localizer fMRI experiment, diffusion tensor imaging, and resting state fMRI (data not analyzed herein). Fifty-five neurotypical adults participated in the same MRI protocol as part of a separate project investigating object representations in the ventral visual pathway (29 females; mean age, 22.27 y; SD 6.03 y). Neurotypical participants had no history of psychiatric illness or neurologic injury and were right-hand dominant (established with the Edinburgh handedness questionnaire^59^. All participants had normal or corrected-to normal vision, took part in the study in exchange for payment, and gave written informed consent in accordance with the University of Rochester Research Subjects Review Board.

### Object Processing Category Localizer fMRI Experiment

fMRI data collection occurred over an 8-year period. Stimulus presentation was controlled with ‘A Simple Framework’^60^ using the Psychophysics Toolbox^61^, with PsychoPy^62^, or with custom stimulus presentation software. Five of the 114 participants with brain lesions data were removed due to technical issues with stimulus presentation software. Participants viewed the stimuli binocularly through a mirror attached to the head coil adjusted to allow foveal viewing of a back-projected monitor (spatial resolution = 1400 × 1050 pixels; temporal resolution = 120 Hz). Each MRI scan consisted of 2 to 8 runs of the object processing category localizer experiment.

Several updates were introduced over the years to the experiment, resulting in three versions: In version A of the category localizer, participants viewed scrambled and intact images of manipulable objects, animals, faces, and places in a miniblock design.^63–67^ There were 12 grayscale photographs per category and 8 exemplars of each stimulus, resulting in 96 images per category, and 384 total images. Phase-scrambled versions of the stimuli were created to serve as a baseline condition. Within each miniblock, 12 stimuli from the same category were presented for 1000 ms each (0 ms interstimulus interval), and 12-second fixation periods were presented between miniblocks. Within each run, 8 miniblocks of intact images and 4 miniblocks of phase-scrambled stimuli were presented with the constraint that a category of objects did not repeat across two successive miniblocks.

In version B, grayscale photographs of manipulable objects, animals, fearful faces, faces with neutral affect, body parts, famous places, and words were used. There were 64 exemplars of fearful faces, neutral faces, and words; for all other categories there were 8 exemplars of each stimulus, resulting 64 images per category, and 448 total images. Within each miniblock, 8 stimuli from the same category were presented for 1000 ms each (0 ms interstimulus interval), and 8-second fixation periods were presented between miniblocks. Within each run, 12 miniblocks of intact images and 2 miniblocks of phase-scrambled stimuli were presented with the constraint that a category of objects did not repeat across two successive miniblocks.

In version C, grayscale and color photographs of single-handed manipulable objects (e.g., hammer), two-handed manipulable objects (e.g., axe), household objects (e.g., book), animals, fearful faces, faces with neutral affect, famous, body parts, famous places, non-famous places, fruits and vegetables, surface textures (e.g., leather), words, and numbers were used. There were 48 exemplars of fearful faces, neutral faces, non-famous places, words, and numbers; for all other categories, there were 6 items and 8 exemplars, resulting in 48 images per category, and 720 total images. Within each miniblock, 6 stimuli from the same category were presented for 1800 ms each (0 ms interstimulus interval). Fixation periods ranging from 4 to 12 seconds (8 seconds on average) were presented between miniblocks. Within each run, 15 miniblocks of intact images and 1 miniblock of phase-scrambled stimuli were presented with the constraint that a category of objects did not repeat across two successive miniblocks.

### MRI Parameters

Whole brain MRI was conducted on a 3-Tesla Siemens MAGNETOM Trio scanner with a 32- or 64-channel head coil at the University of Rochester. High-resolution structural T1 contrast images were acquired using a magnetization-prepared rapid gradient echo pulse sequence at the start of each participant’s scanning session (TR = 2530, TE = 3.44ms, flip angle = 7°, FOV = 256mm, matrix = 256 × 256, 192 1 x 1 x 1mm sagittal left-to-right slices). An echo-planar imaging pulse sequence was used for T2* contrast. In the first 3 participants with lesions and 55 neurotypical participants, the acquisition parameters were: TR = 2000ms, TE = 30ms, flip angle = 90°, FOV = 256 × 256 mm, matrix = 64 × 64, 30 inferior-to-superior axial slices, voxel size = 4 x 4 x 4 mm. In the next 40 participants with lesions, the acquisition parameters were: TR = 2200ms, TE = 30ms, flip angle = 90°, FOV = 256 × 256 mm, matrix = 64 × 64, 33 inferior-to-superior axial slices, voxel size = 4 x 4 x 4mm. In the remaining 66 participants with lesions, the acquisition parameters were upgraded to a multi-band sequence: TR = 2200ms, TE = 30ms, flip angle = 70°, FOV = 256 × 256 mm, matrix = 128 × 128, 90 inferior-to-superior axial slices, voxel size = 2 x 2 x 2 mm. Across all participants, the first 4 volumes of each run were discarded to allow for signal equilibration (4 volumes dropped prior to image acquisition).

Diffusion MRI was acquired using a single-shot echo-planar sequence (60 diffusion directions with b = 1000 s/mm^2^, 10 images with b = 0 s/mm^2^, TR = 8900 msec, TE = 86 msec, FOV = 256 x 256mm, matrix = 128 x 128, voxel size = 2 x 2 x 2 mm, 70 inferior-to-superior axial slices; anterior-to-posterior phase encoding). A double-echo gradient echo field map sequence (echo time difference = 2.46 msec, EPI dwell time = 0.75 msec) was acquired with the same resolution as the DTI sequence to correct for distortion caused by B0 inhomogeneity.

### Lesion Identification

Lesions were identified by segmenting healthy tissue from the damaged tissue using the native T1 image. Lesioned voxels consisting of both grey and white matter were assigned a value of 1, and preserved voxels were assigned a value of 0 using ITK-SNAP. All participants provided written consent for the research team to access clinical MRI acquired as part of their clinical care. Lesion drawings were reviewed by a neuro-radiologist (co-author S.P. Meyers), who was at the time naïve to the aims of the project, and who had access to T1, T2, and FLAIR imaging (with and without contrast). Lesion renderings were inspected by the neuro-radiologist and corrected by study team members when they did not accurately capture the boundaries of the lesion. Lesion renderings in the native T1 space were entered as cost function masks^68^ during anatomical and functional data pre-processing. This feature ensures that the anatomical data could be normalized to Montreal Neurological Institute (MNI) space while minimizing error introduced by a lesion. The corrected lesion drawings were normalized to MNI space using the transformation matrix that normalized the native T1 into MNI space (using the ANTs toolbox). Finally, the accuracy of the lesions in MNI space were reviewed and confirmed, using anatomical landmarks as reference to ensure no distortions were introduced during normalization.^29,69^

### Anatomical MRI Pre-processing

Each participant’s T1-weighted images was corrected for intensity non-uniformity with ‘N4BiasFieldCorrection’ (ANTs), and was used as the T1-reference. The T1-reference was skull-stripped with the ‘antsBrainExtraction.sh’ workflow using OASIS30ANTs as target template. Brain tissue segmentation of cerebrospinal fluid, white matter, and gray matter was performed on the brain-extracted T1 using FSL’s ‘fast’ tool. Volume-based spatial normalization to standard space (MNI152NLin2009cAsym) was performed through nonlinear registration with ‘antsRegistration’, using brain extracted versions of both T1 reference and the T1 template. The ICBM 152 Nonlinear Asymmetrical template version 2009c was selected for spatial normalization.

### Functional MRI Pre-processing

For each run of fMRI data, a reference volume and its skull-stripped version were generated using fMRIPrep.^70^ Head-motion parameters with respect to the BOLD reference (transformation matrices, and six corresponding rotation and translation parameters) were estimated before spatiotemporal filtering using ‘MCFLIRT’ from FSL. BOLD runs were slice-time corrected using ‘3dTshift’ from AFNI. The BOLD time-series were then resampled onto their original, native space by applying the transforms to correct for head-motion. The BOLD reference was then co-registered to the T1 reference using ‘mri_coreg’ (FreeSurfer) followed by FSL’s FLIRT tool with the boundary-based registration cost-function. Co-registration was configured with twelve degrees of freedom to account for distortions remaining in the BOLD reference. The BOLD time-series were resampled into standard space, generating a pre-processed BOLD run in MNI152NLin2009cAsym space. Following pre-processing, fMRI and T1 anatomical data were analyzed with the BrainVoyager software package (Version 22.4) and in-house scripts drawing on the NeuroElf toolbox in MATLAB. Functional data underwent spatial smoothing (6 mm FWHM) and temporal high-pass filtering (cutoff: 2 cycles per time course per run) after voxels were interpolated to 3 x 3 x 3 mm^3^. For all participants, a general linear model was used to fit beta estimates to the experimental events of interest. Experimental events were convolved with a standard 2-gamma hemodynamic response function. The first derivatives of 3D motion correction from each run were added to all models as regressors of no interest to attract variance attributable to head movement.

### Diffusion MRI Pre-processing

FSL’s BET tool skull-stripped each participant’s diffusion and T1 images, as well as the fieldmap magnitude image. The B0 image was stripped from the diffusion weighted image, and the fieldmap was prepared using FSL’s prepare fieldmap tool. Smoothing and regularization was performed using FSL’s fugue tool and a 3D Gaussian smoothing was applied (sigma = 4 mm). The magnitude image was warped based on this smoothing, with *y* as the warp direction. Eddy current correction was performed using FSL’s eddy_correct tool, which takes each volume of the diffusion-weighted image and registers it to the b0 image to correct for both eddy currents and motion. The deformed magnitude image was registered to the b0 image using FSL’s FLIRT tool. The resulting transformation matrix was then applied to the prepared fieldmap. Lastly, the diffusion-weighted image was undistorted using the registered fieldmap with FSL’s fugue tool. Intensity correction was also applied to this unwarping. This procedure was applied to all but 3 participant datasets. Susceptibility artifact in the 3 remaining datasets was corrected using FSL’s TOPUP tool. The outputs of TOPUP served as inputs to FSL’s eddy_correct tool.

### ROI Definition: Manipulable Object Preferring Left Ventral Temporal Cortex

Strict independence of the criteria for voxel definition and test was applied throughout to avoid ‘double dipping’ ^71^ in localizing voxels preferring manipulable objects. The whole-brain contrast of ‘Tools > [Animals, Faces, and Places]’ (equally weighted) was computed at the participant level. If this contrast did not elicit a response below a False Discovery Rate (FDR) corrected value (*q* < .05), the threshold was relaxed to *p* < 0.05, uncorrected. We began by identifying the peak voxel in left VTC using all runs of data; a spherical ROI 3 mm in diameter was then drawn on the peak (see Figure 1A, C). Leave-one-out cross-validation was used to identify the peak voxel within that sphere using *N* – 1 runs of data. A new spherical ROI 3 mm in diameter was then drawn on the N-1 defined peak voxel prior to extracting *t*-values for the contrast of ‘Tools > [Animals, Faces, and Places]’ (equally weighted) using the left-out *N*th run. VTC voxels that preferred manipulable objects defined using *N*-1 runs were used to extract the response to place stimuli with all runs, using the contrast of ‘Places > [Animals, Faces, and Tools]’ (equally weighted). After iterating this procedure *N* times, we averaged across data folds to obtain a mean response for manipulable objects and places for each participant, which served as the independent variables in a Voxel-based Lesion Activity Mapping (VLAM) analysis. This localization scheme identified voxels preferring manipulable objects in left VTC of 95 of the 109 participants with brain lesions and 52 of the 55 neurotypical participants (see Supplemental Table 1 for values).

**Figure 1.**
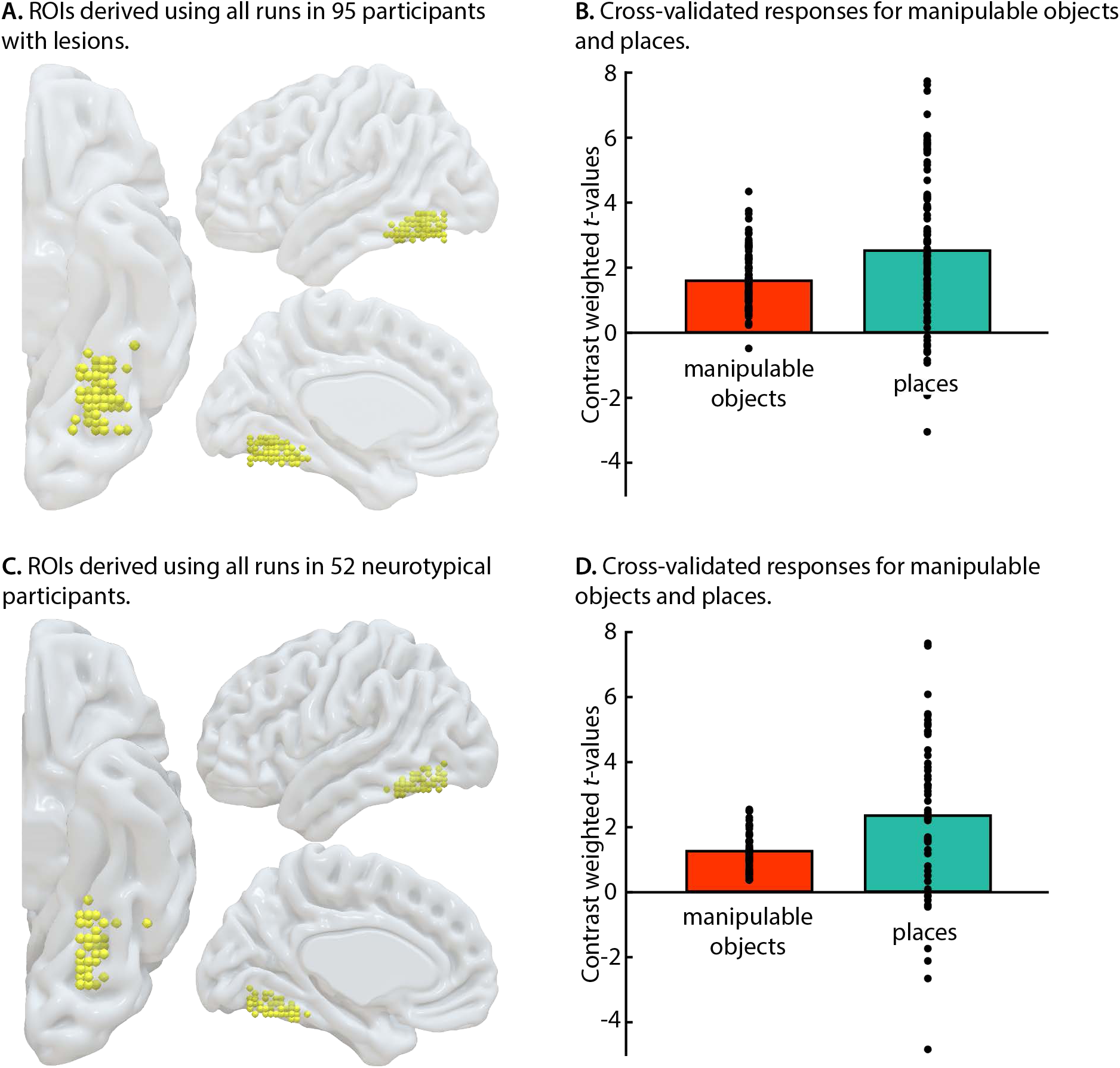
Preferences for manipulable objects and places in left VTC. **A., C.** Spheres 3 mm in diameter represent the participant-specific manipulable object-preferring voxels localized using all runs. **B., D.** A leave-one-out cross-validation approach used a spherical region-of-interest (ROI) localized using N-1 runs of data to extract manipulable object preferences from the *N*th run using the contrast of ‘Tools > [Animals, Faces, and Places]’ (equally weighted). This procedure was iterated N times and averaged across N data folds to derive a cross-validated *t*-value for manipulable objects. The same ROIs were used to obtain cross-validated *t*-value for place stimuli using the contrast of ‘Places > [Animals, Faces, and Tools]’ (equally weighted).

### Voxel-based Lesion Activity Mapping

VLAM uses functional neural responses in an ROI to predict lesion overlap throughout the brain. VLAM was performed using the SVR-LSM toolbox using a univariate model^72^. Prior to the VLAM analysis, each participant’s lesion mask was interpolated from 1×1×1 mm to 2×2×2 mm to reduce the number of multiple comparisons performed. Voxels lesioned in at least 10% of participants were analyzed. We controlled for variability in lesion volume using the ‘Regress Both’ option, which regresses variability in total lesion volume from fMRI responses and from every lesioned voxel entered in the analysis. Voxelwise statistical significance was determined using a permutation analysis in which the fMRI responses were randomly assigned to a lesion map, and the same procedure as described above was iterated 10,000 times. Voxelwise z-scores were computed for the true data in relation to the mean and standard deviation of voxel-by-voxel null distributions; the resulting z-score map was set to a threshold of z < −1.65 (*p* < .05, one-tailed) to determine chance-level likelihood of identifying a significant effect in each voxel. If the resulting voxelwise clusters did not survive permutation analysis, we removed clusters with fewer than 500 contiguous voxels before interpreting the results^73–77^. The analysis was then re-run using place preferences derived from the participant-defined left VTC ROIs. The Harvard-Oxford cortical and subcortical probabilistic atlases assessed the location of significant voxels identified using VLAM.

### Preparation of diffusion MRI data for connectome-based lesion activity mapping analyses

Seventy-five of the 95 participants in whom we could localize voxels preferring manipulable objects in the left VTC took part in the 60-direction diffusion MRI protocol, and were thus included in the white matter fiber tracking analysis. Of the 20 remaining participants, 1 did not take part in diffusion MRI, and 19 took part in a 258-direction diffusion spectrum imaging protocol (data not analyzed herein). Fractional anisotropy (FA) along each voxel of the eddy corrected diffusion MRI data was modeled within DSI_Studio. Each participant’s FA map was then co-registered to the ICBM152 template, and entered into a connectome-wide analysis. The connectome-wide analysis uses whole-brain multiple regression with Spearman correlation to relate variance in voxelwise FA to variance in manipulable object preferences in the left VTC after controlling for total lesion volume. Voxelwise correlations exceeding a *t*-statistic of 3.00 were entered into a deterministic tractography analysis with the VLAM-identified lesion site entered as an ROI (see Figure 3, red-to-yellow). The fiber tracts identified meet the dual criteria of accounting for a significant amount of variance in manipulable object responses in the left VTC, and exhibit structural connectivity to the VLAM-identified lesion site. A 10,000 iteration permutation analysis was then performed in which the fMRI responses were randomly assigned to FA maps. For each permutation, multiple regression was performed and significant voxels were entered into an identical deterministic tractography analysis with the VLAM-identified lesion site as the ROI. The length of fiber tracts identified in the permutation analysis were compared against the length of fiber tracts in the non-permuted analysis to estimate the false discovery rate (*q* < .05) using a length threshold of 15. Fiber tracts that survived FDR correction (i.e., fibers that account for variance in manipulable object responses in the left VTC and that are significantly greater than 15 voxels in length) are reported herein.

## Results

### Analysis of preferences for manipulable objects and places in the left ventral temporal cortex

Regions of left VTC that preferred manipulable objects were able to be localized using a leave-one-out approach in 95 of the participants with brain lesions (88%; Figure 1A) and 52 of the 55 neurotypical participants (96%; Figure 1C). We first sought to establish that neural preferences in the left VTC for manipulable objects and places were comparable across the groups. To that end, we conducted a mixed ANOVA with the between-subject factor of Group and the within-subject factor of Category. There was a main effect of Category (*F*(1, 145) = 20.56, p < .001, ƞ² = 0.12), as responses to places were stronger than responses to manipulable objects. This finding replicates prior studies^29,49,52^, and shows that non-manipulable stimuli (e.g., places, scenes, houses) drive, if anything, stronger neural responses in VTC voxels that also show differentially high functional responses for manipulable objects. There was no effect of Group nor an interaction between Group and Category (*F* < 1), indicating equivalently larger functional responses for places compared to manipulable objects in participants with lesions (Figure 1B) and in neurotypical participants (Figure 1D).

### aIPS and SMG lesions dampen neural responses for manipulable objects in left VTC

In the VLAM analyses performed here, we entered the contrast-weighted *t*-values for manipulable objects in left VTC as the independent variable (Figure 1B), and tested where lesion presence (dependent variable) was inversely related to the amplitude of responses. The VLAM analysis showed that lesion presence in the left inferior parietal lobule was associated with lower amplitude neural responses to manipulable objects in left VTC. This included the SMG, extending into the aIPS and parietal operculum and posteriorly into the angular gyrus. The analysis also identified the posterior middle temporal gyrus as a region where lesions were associated with reduced neural responses for manipulable objects in VTC (Figure 2A, red). This result extends a prior demonstration with 35 participants with lesions^29^ (see Table 1A for voxel-level statistics).

**Figure 2.**
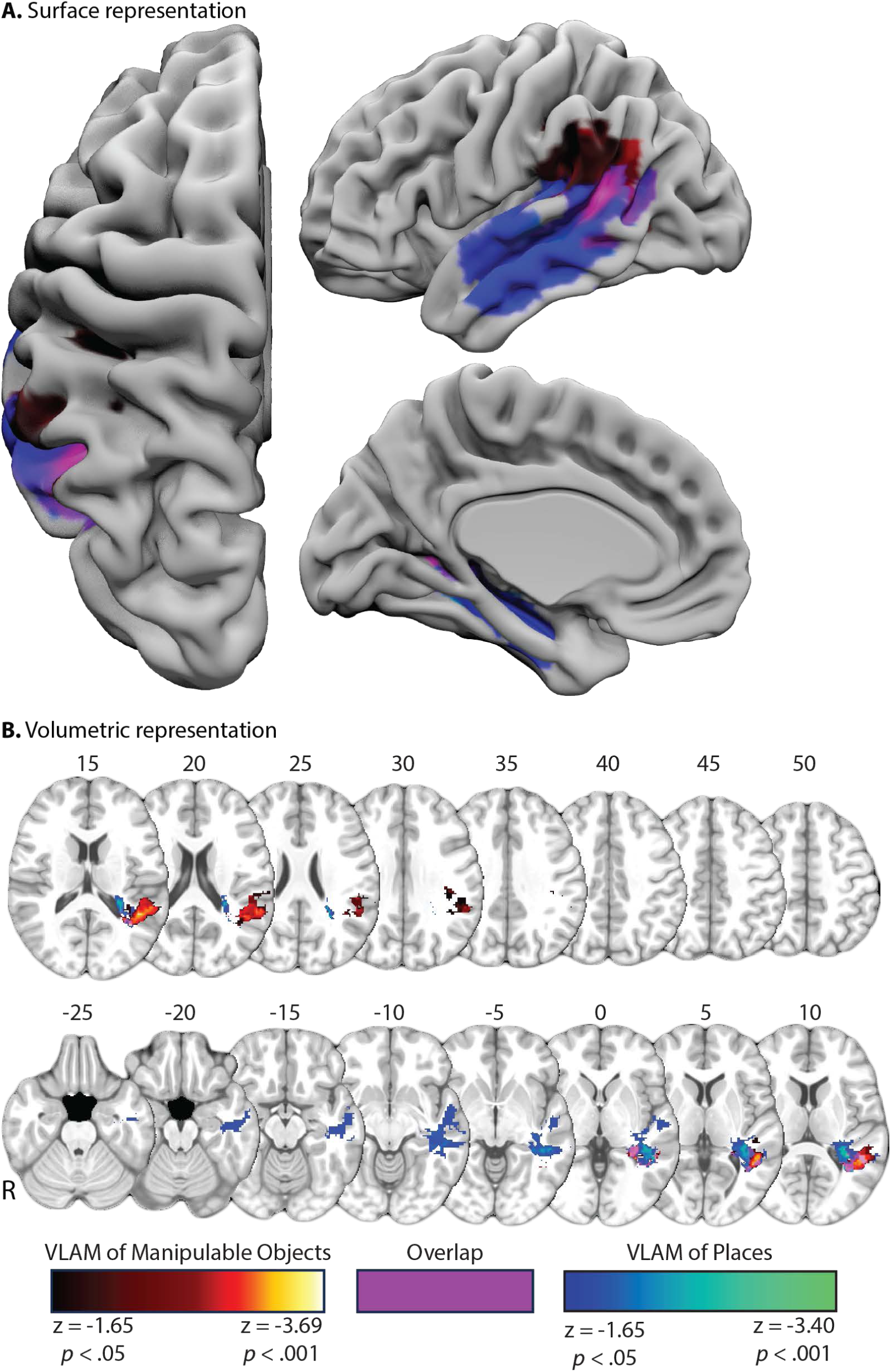
VLAM analysis of manipulable objects and places in left VTC. **A.** Reduced preferences for manipulable objects in left VTC were associated with lesion presence in the left SMG, aIPS, parietal operculum, angular gyrus, and the posterior middle temporal gyrus (red-to-yellow). An overlapping lesion site in the posterior temporal lobe was also associated with reduced place preferences (magenta), whereas lesions to anterior portions of the temporal lobe, thalamus, insula, hippocampus, and posterior fusiform gyrus were uniquely associated with place preferences in left VTC (blue-to-green).

When repeating the analysis with place preferences as the independent variable, we identified a site in the lateral temporal lobe, including the middle temporal and superior temporal gyri, and a portion of white matter undercutting the parietal operculum (Figure 2, blue). A second lesion site included the thalamus, insula, and hippocampus (see Table 1B for voxel-level statistics). Critically, the voxels identified in the VLAM analysis of places did not overlap with the parietal lesion site identified in the VLAM analysis of manipulable objects. The overlap between the two analyses was principally in the superior and middle temporal gyri (Figure 2, purple).

### Direct comparison between VLAM analyses for manipulable objects and places

Next, we tested whether the VLAM result for manipulable objects that identified the SMG and aIPS was statistically stronger than the corresponding VLAM result for places, using a permutation test to estimate the likelihood of the observed differences. Figure 3 (red-to-yellow) shows the voxels for which lesion presence *differentially* affected functional responses to manipulable objects, compared to places, in the left VTC. This analysis identifies the same left SMG and aIPS subregions identified in the VLAM analysis of manipulable objects in Figure 2. A smaller cluster of voxels were identified in the left parietal operculum, and the left posterior middle temporal gyrus (see Table 2A for voxel-level statistics). These results further reinforce that lesions to the left SMG and aIPS differentially affect neural preferences to manipulable objects in left VTC. Critically, this result is obtained when using the most stringent baseline possible—place stimuli, which elicit, if anything, stronger responses in the relevant VTC regions compared to manipulable objects. In other words, because overall neural response to manipulable objects are lower than neural responses to places in the VTC^49,52^, this represents the strongest possible test of the specificity of the VLAM results to manipulable objects.

**Figure 3.**
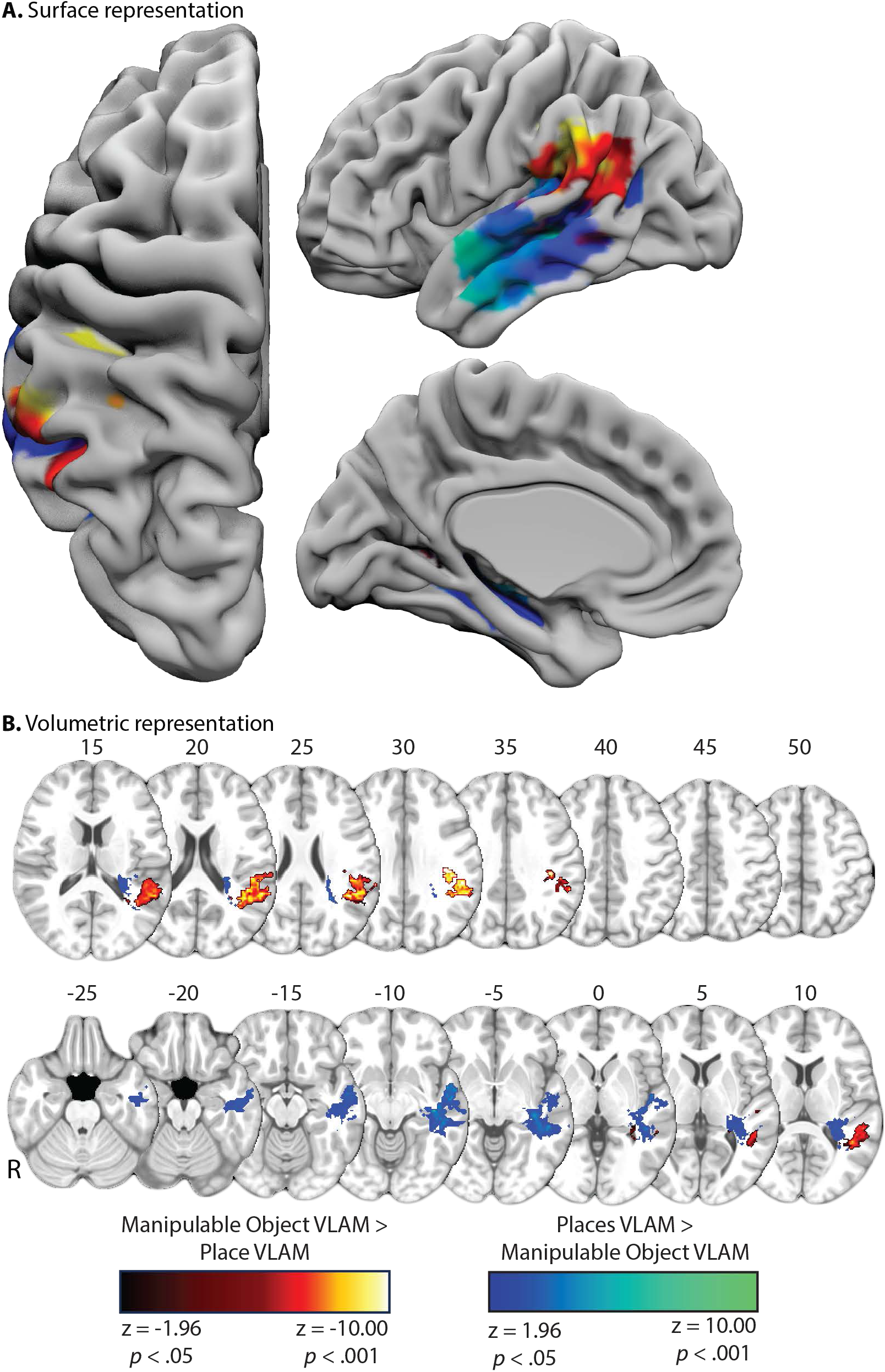
Voxelwise comparison between VLAM analyses of manipulable objects and places in the left VTC. Values in each VLAM map in Figure 2 are z-scores; thus, we performed a subtraction analysis in each voxel to quantify the difference between the result for manipulable objects subtracting out the result for places. A 1,000 iteration permutation analysis followed, in which the VLAM maps were scrambled prior to subtracting the place VLAM map from the manipulable object VLAM map. The voxelwise difference scores of the true data were then z-scored against the mean permutation-derived voxelwise difference scores. The voxelwise comparison analysis identifies the left SMG, aIPS, posterior middle temporal gyrus and underlying white matter as a VLAM-identified lesion site that was significantly greater for manipulable objects. By contrast, lesions to the anterior inferior, middle, and superior temporal gyri, as well as the thalamus, insula, hippocampus, and posterior fusiform gyrus, were associated with a stronger VLAM result for places.

By contrast, Figure 3 (blue-to-green) shows that lesions to the anterior inferior, middle, and superior temporal gyri, the thalamus, insula, and hippocampus were associated with a stronger VLAM result for places (see Table 2B for voxel-level statistics). This extends a previous demonstration that lesions to ventral and lateral portions of the left temporal lobe were associated with reduced responses for places in left VTC^29^.

### Connectome-based lesion activity-mapping of manipulable objects and places

We then tested which white matter pathways, involved in connectivity between the left inferior parietal lobule and left VTC, are linked to the disruption of VTC functional responses caused by inferior parietal lesions. Connectometry is a technique in which variance in fractional anisotropy (FA) throughout the brain is predicted by an independent variable^78^, in this case, contrast-weighted *t*-values for manipulable object preferences in left VTC (for other variants of this method, see ^79,80^). Variability in total lesion volume was regressed from the amplitude of manipulable object preferences prior to conducting the analysis. This approach defines, whole-brain, voxels for which variance across patient participants in FA values is related to variance in functional neural responses for manipulable objects in the left VTC. Once identified, those voxels were entered into a deterministic tractography analysis, using the inferior parietal and lateral temporal lobe voxels identified in voxelwise comparison of VLAM effects (Figure 3, red). The fiber tracts identified in this manner meet the dual criteria of exhibiting: ***i)*** a significant association with neural responses to manipulable objects in left VTC, and ***ii)*** support structural connectivity to the VLAM-identified lesion site in the inferior parietal and lateral temporal lobe sites. A second, parallel, connectometry analysis was run with place preferences in left VTC as the independent variable. Voxels identified were entered into a deterministic tractography analysis using the temporal lobe lesion site identified in the voxelwise comparison of VLAM results (Figure 3, blue).

The connectometry analysis of manipulable objects identified a fiber bundle that connects the left VTC to the left inferior parietal lobule principally via the arcuate fasciculus (Figure 4, yellow). Although the left inferior fronto-occipital fasciculus, the left inferior longitudinal fasciculus, and the left middle longitudinal fasciculus were identified as fibers connecting the left VTC to the left inferior parietal lobule in the context of manipulable objects (Figure 4A, red), those tracts overlapped with the white matter pathways identified in the connectometry analysis of places (Figure 4, blue; see Table 3). When color-coding the fiber tracts identified in the connectometry analysis of manipulable objects, there is convergence between the VLAM-identified aIPS and SMG (Figure 4B, red) and the connectometry-identified left arcuate fasciculus (Figure 4B, yellow; axial slice Z = 30). Inferior to that lesion site is a second area in which the VLAM-identified lesion site in the posterior middle temporal gyrus overlaps with the connectometry-identified inferior fronto-occipital fasciculus (Figure 4B, green), inferior longitudinal fasciculus (Figure 4B, blue), and middle longitudinal fasciculus (Figure 4B, purple). Although a small portion of these ventral longitudinal fibers were identified in the connectometry analysis of places, the left arcuate fasciculus was specific to manipulable objects (see Table 3).

**Figure 4.**
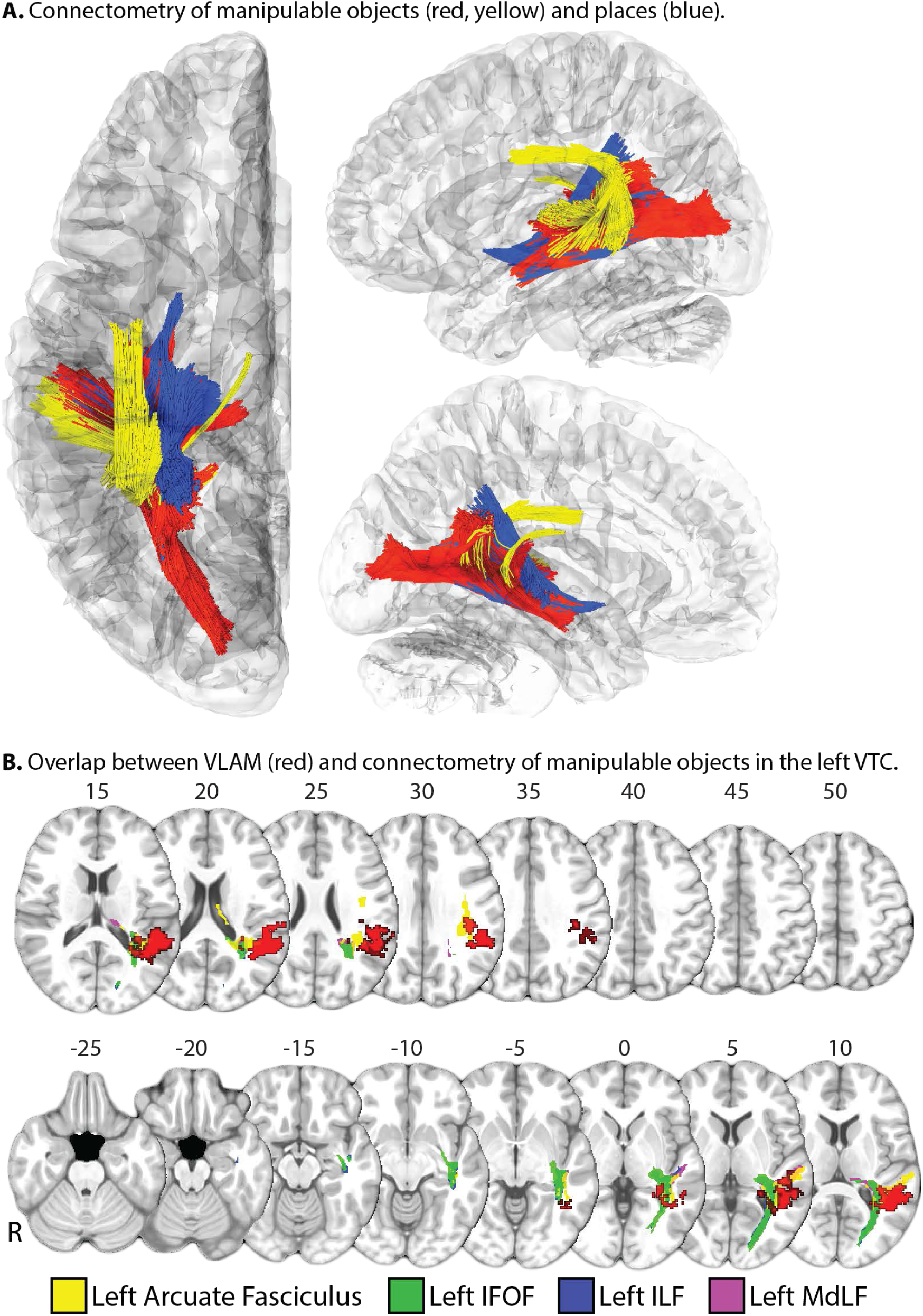
Connectome-based lesion-activity mapping in the left VTC. **(A)** The connectometry analysis of manipulable objects (red, yellow) and places (blue) identified overlapping fibers in the left inferior fronto-occipital fasciculus, the left inferior longitudinal fasciculus, and the left middle longitudinal fasciculus. By contrast, the arcuate fasciculus (yellow) was identified as a white matter pathway in which reduced FA was associated with reduced responses for manipulable objects in left VTC. **(B)** There is convergence between the VLAM-identified aIPS and SMG lesion site (red) and the connectometry-identified left arcuate fasciculus (yellow; axial slice Z = 30). Inferior to that is a lesion site in the posterior middle temporal gyrus that overlaps with the inferior fronto-occipital fasciculus (green), inferior longitudinal fasciculus (blue), and middle longitudinal fasciculus (purple).

## General Discussion

We have reported causal evidence for an integrated network that allows object representations in the temporal lobe to interface with action representations in the parietal lobe: Lesions involving the left aIPS and SMG (Figure 2) reduce functional neural responses to manipulable objects in left VTC. We also identified the white matter tracts in which reduced fractional anisotropy was associated with reductions in manipulable object preferences in the left VTC. These tracts were composed of ventral longitudinal fibers, including the inferior fronto-occipital fasciculus, the inferior longitudinal fasciculus, and the middle longitudinal fasciculus, and were structurally connected to the left inferior parietal via the left arcuate fasciculus (Figure 4). Of those pathways, the descending portion of the left arcuate fasciculus was specifically related to reductions in functional neural responses to manipulable objects, compared to places, in VTC. Collectively, our results demonstrate that lesions to this network interfacing temporal lobe object representations with parietal action representations causes a category-specific disruption in neural responses VTC regions distal to the site of injury. This general phenomenon was first described by Price and colleagues (2001), who referred to it as a type of dynamic diaschisis (see also ^82,83^).

The VLAM analysis of manipulable object responses in VTC also identified the lateral temporal lobe in the vicinity of the posterior middle temporal gyrus, which suggests that downstream conceptual processing may affect processing at earlier stages of the visual hierarchy^84,85^. The results of the connectometry analysis support this position, as reduced structural integrity of the left arcuate fasciculus, a tract mediating structural connectivity between the left inferior parietal lobule and left posterior middle temporal gyrus^53^, was associated with reduced manipulable object preferences in left VTC. It is known that the dorsal visual pathway has access to perceptual representations^86,87^ such as the shape and location of the object with respect to the body^88^, and whether an object is elongated^89,90^. These representations provide an initial grasp target, but ultimately, visuomotor processing requires inputs about the conceptual and perceptual attributes of the object to maximize end state comfort^11^. Neuropsychological evidence indicates that that the dorsal stream can process, independent of the ventral stream, rudimentary information about grasp targets^6^. Compelling observations from blindsight have demonstrated preserved visuomotor abilities to visual targets presented in the hemianopic visual field in individuals with post-thalamic lesions. Specifically, individuals exhibiting ‘action blindsight’ ^91^ have demonstrated intact grip scaling and wrist orienting for hand actions^92,93^, and the ability to navigate obstacles^94^. Those observations demonstrate that the dorsal visual pathway receives substantial (and for some tasks, *sufficient*) input from extra-geniculostratiate pathways about the volumetry, orientation, and location of objects in peripersonal space^95–97^.

Our proposal is that an initial low-resolution representation of a visual stimulus as an action target (via the dorsal stream) triggers finer-grained analyses integrating material properties (ventral and medial temporal cortex) with conceptual representations (posterior middle temporal gyrus) into the action plan. Ultimately, actions toward objects are based on an understanding of what will be done with the object in order to satisfy behavioral goals. For instance, if a glass is upside down on the counter, how it is grasped will depend on whether the goal is to put it in the cupboard upside down or to take a drink of water. This analysis takes place prior to and in parallel with processing within the ventral visual pathway^8,98–100^, and may serve to bias processing within the ventral visual pathway to extract visual form, surface texture, and materials properties to facilitate functional grasping and object manipulation.

Material properties such as surface texture and object weight are processed in medial and ventral temporal cortex^43,44^, which overlaps with the voxels exhibiting preferences for manipulable objects in left VTC. Integration of object material properties into the action plan is a key computational goal that could be achieved by the parietal-to-VTC connectivity we have described. This dorsal-to-ventral integration model is aligned with the approach emphasized by Bar and colleagues^101,102^, who showed that the orbitofrontal cortex uses a magnocellularly driven low-acuity representation of visual stimuli to guide subsequent fine-grained visual processes in the ventral visual pathway^103–105^.

The connectometry analysis of manipulable objects identified the left inferior fronto-occipital fasciculus, the left inferior longitudinal fasciculus, and the left middle longitudinal fasciculus. Whereas structural integrity of the left inferior fronto-occipital fasciculus has been found to predict tool use accuracy in patients with brain lesions^58^, the inferior and middle longitudinal fasciculi have not been implicated in prior studies of object-directed action. One possibility is that these pathways integrate processing across a hierarchy of visual regions to support both feedforward and feedback processing, irrespective of the stimulus content. Lesions disrupting these processes can reduce functional responses for manipulable objects and places in the VTC, as we have observed here. By contrast, the descending portion of the left arcuate fasciculus was identified as a white matter tract in which reduced FA was associated with manipulable object preferences in left VTC. Human white matter dissection studies have shown that a portion of the posterior descending segment of the left arcuate fasciculus terminates in the posterior middle temporal gyrus^106^, which aligns with the segment associated with manipulable object responses in the VTC (Figure 4, yellow). This finding further reinforces the notion that neural responses in the VTC are modulated both by processes that result from a bottom up analysis of the visual input, and by inputs from parietal action system, mediated by structural connectivity via the arcuate fasciculus.

The proposal that lesions to praxis representations in the left inferior parietal lobule disrupt neural responses for manipulable objects in left VTC suggests that the motor computations underlying skilled action provide modulatory feedback to analysis of the material properties of objects in the VTC. One articulation of this framework is that activation of praxis representations includes predictions about how an action should look and feel when produced, which incorporates real-time feedback of material properties in the VTC to update the current and future state of the hand and arm during object-directed action^12,18,29^. We have provided causal evidence that lesions to the left inferior parietal lobule and the descending portion of the left arcuate fasciculus cause a category-specific disruption in processing in VTC. More broadly, these findings suggest that distal structural connectivity is critical in shaping the topography by semantic category within the ventral visual pathway. Future studies leveraging causal evidence from white matter dissection mapping will be crucial to advance understanding of the structural basis of functional diaschisis in the human brain.

## Data availability

The whole-brain maps of contrast-weighted t-values, lesions in MNI space, ROIs localized, and analysis code will be made available via a publicly accessible repository upon publication of the manuscript. Participants did not consent to public archival of the MRI data. Researchers interested in accessing the raw MRI data should contact the corresponding author to establish a data use agreement.

## Acknowledgments

This research was supported by the University of Rochester CTSA awards NIH UL TR002001 and NIH KL2 TR001999 from the National Center for Advancing Translational Sciences, and from the Harry W. Fischer fund within the Department of Imaging Sciences at the University of Rochester Medical Center to F.E.G. This research was also supported by NIH grants R21NS076176 and R01NS089069 and NSF grant BCS-1 349 042 to B.Z.M., by a core grant to the Center for Visual Science (P30 EY001319), and by funding support to the Department of Neurosurgery at the University of Rochester by Norman and Arlene Leenhouts.

