## Supplemental Online Materials for "Visual processing of manipulable objects in the ventral stream is modulated by inputs from parietal action systems"

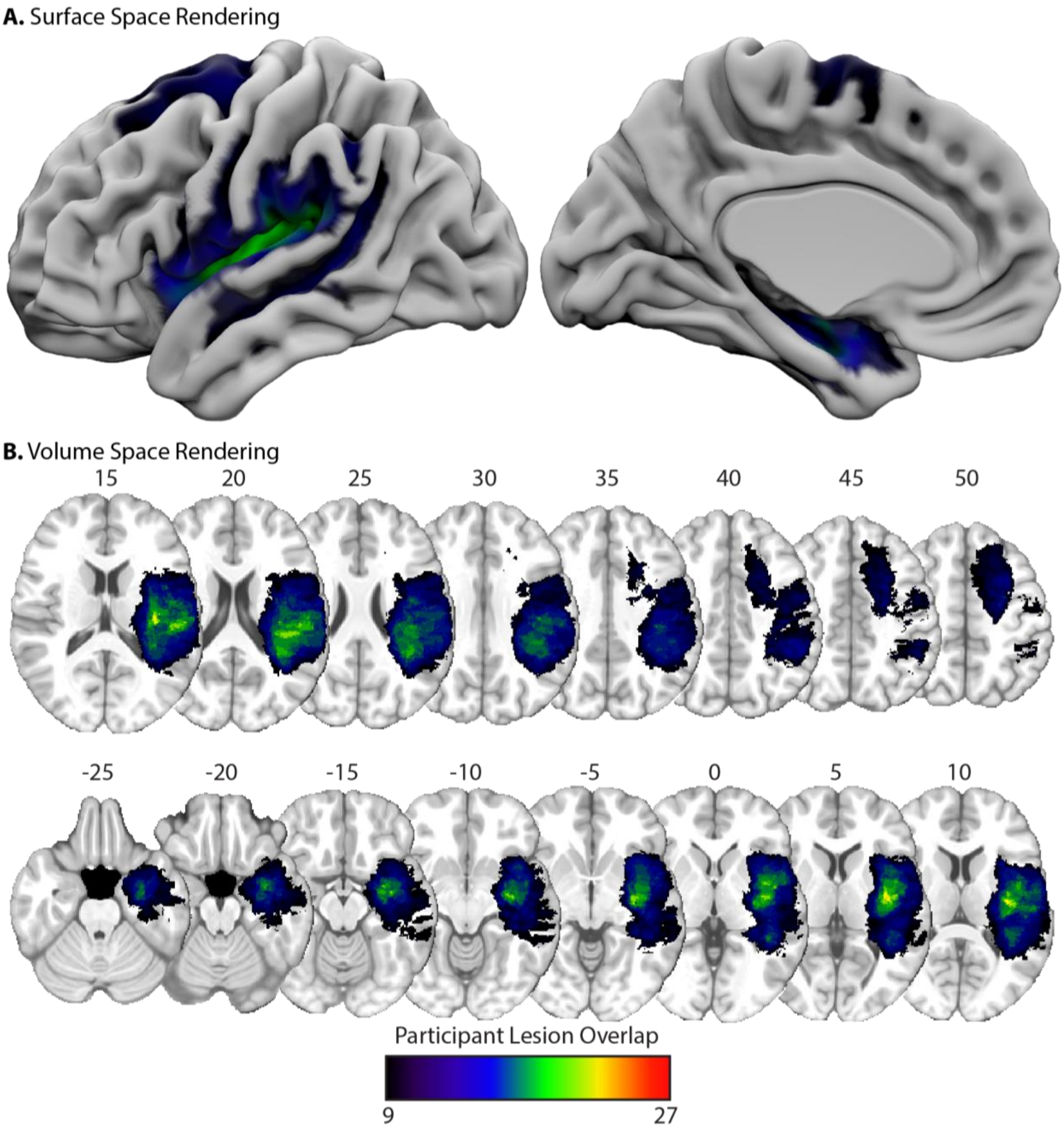


**Supplemental Figure 1. Voxelwise lesion overlap among participants with a left hemisphere lesion.** (A) Projected on the cortical surface is lesion overlap among participants for whom we could localize voxels preferring manipulable objects in the left ventral temporal cortex. Warmer colors indicate greater lesion overlap among participants. (B) The same map is projected in volume space (Z-coordinate provided). The areas of greatest lesion overlap include the left anterior temporal lobe, hippocampus, insula, pre- and post-central gyri, the superior frontal gyrus, the posterior superior temporal gyrus, and the supramarginal gyrus.

**Supplemental Table 1.** Demographic information for each participant. *Abbreviations*. Dysembryoplastic Neuroepithelial Tumor, DNET; GBM, glioblastoma multiforme; AVM, arteriovenous malformation; PLNTY, polymorphous low grade neuroepithelial tumor of the young.

**A.** Participants with brain lesions

| Cumulative Participant ID | Age | Diagnosis | Gender | Lesion Volume | Hemisphere | T>AFP *t*-values | P>AFT *t*-value |
|---|---|---|---|---|---|---|---|
| 1 | 52 | Adenocarcinoma | Female | 3018 | Left | 1.59 | -0.85 |
| 2 | 73 | GBM | Male | 11349 | Left | Could not localize ROIs | |
| 3 | 58 | Adenocarcinoma | Female | 85097 | Left | 1.14 | 2.55 |
| 4 | 58 | Astrocytoma | Female | 363396 | Left | 1.04 | 3.79 |
| 5 | 69 | Oligodendroglioma | Female | 99764 | Left | 3.00 | 0.45 |
| 6 | 47 | DNET | Female | 34086 | Left | -0.49 | 1.19 |
| 7 | 69 | GBM | Male | 31102 | Left | Could not localize ROIs | |
| 8 | 26 | Astrocytoma | Female | 39900 | Right | Could not localize ROIs | |
| 9 | 47 | Oligodendroglioma | Male | 99956 | Left | 1.21 | 3.77 |
| 10 | 30 | Astrocytoma | Male | 57742 | Right | 0.75 | 1.78 |
| 11 | 29 | Astrocytoma | Male | 20878 | Left | Could not localize ROIs | |
| 12 | 58 | GBM | Female | 188060 | Left | 1.18 | -3.03 |
| 13 | 57 | GBM | Male | 145300 | Left | 1.53 | 4.11 |
| 14 | 54 | Intracerebral hemorrhage | Male | 130675 | Left | 1.97 | 7.65 |
| 15 | 46 | Oligodendroglioma | Male | 67034 | Left | Could not localize ROIs | |
| 16 | 51 | Astrocytoma | Female | 36524 | Right | 1.54 | 1.10 |
| 17 | 27 | Oligodendroglioma | Male | 21384 | Right | 2.77 | 1.20 |
| 18 | 25 | Cavernous malformation | Female | 3993 | Left | 1.96 | 7.35 |
| 19 | 45 | GBM | Female | 127084 | Left | Data discarded due to technical issue | |
| 20 | 28 | Astrocytoma | Male | 84788 | Left | 1.02 | 3.15 |
| 21 | 45 | GBM | Female | 277109 | Left | 2.83 | 5.47 |
| 22 | 48 | Oligodendroglioma | Male | 153685 | Left | 1.47 | 2.05 |
| 23 | 42 | Astrocytoma | Male | 44270 | Left | 4.29 | -1.92 |
| 24 | 56 | Astrocytoma | Female | 78212 | Right | 1.25 | -0.61 |
| 25 | 71 | GBM | Male | 207337 | Left | 0.67 | 1.84 |

| | | | | | | | |
|---|---|---|---|---|---|---|---|
| 26 | 36 | GBM | Male | 84346 | Right | 1.63 | 4.63 |
| 27 | 37 | Cavernous malformation | Female | 183 | Left | 1.01 | 4.05 |
| 28 | 21 | Cavernous malformation | Female | 1708 | Right | 0.52 | 5.81 |
| 29 | 63 | GBM | Male | 79084 | Left | Data discarded due to technical issue | |
| 30 | 68 | AVM | Female | 50224 | Right | Could not localize the LMFG ROI | |
| 31 | 61 | GBM | Female | 133252 | Right | Could not localize the LMFG ROI | |
| 32 | 43 | AVM | Female | 12456 | Left | Data discarded due to technical issue | |
| 33 | 66 | Astrocytoma | Male | 34734 | Left | 2.26 | 6.64 |
| 34 | 67 | Oligodendroglioma | Male | 42216 | Left | 1.36 | 5.60 |
| 35 | 67 | GBM | Male | 18604 | Left | Could not localize the LMFG ROI | |
| 36 | 28 | GBM | Male | 193209 | Left | 2.33 | -0.25 |
| 37 | 36 | Astrocytoma | Female | 64557 | Left | 3.61 | 3.82 |
| 38 | 49 | GBM | Female | 11801 | Left | 3.70 | 3.79 |
| 39 | 65 | Astrocytoma | Male | 29215 | Right | 3.46 | -0.6 |
| 40 | 55 | Astrocytoma | Female | 15297 | Left | 1.43 | 2.32 |
| 41 | 20 | Astrocytoma | Male | 7570 | Left | 2.20 | 5.87 |
| 42 | 23 | Astrocytoma | Male | 19704 | Left | 2.12 | 2.73 |
| 43 | 57 | Oligodendroglioma | Male | 9308 | Left | 1.16 | -0.37 |
| 44 | 32 | Oligodendroglioma | Male | 19057 | Left | 1.61 | 5.97 |
| 45 | 29 | Astrocytoma | Female | 69361 | Left | Data discarded due to technical issue | |
| 46 | 39 | AVM | Male | 12433 | Left | 0.76 | 2.97 |
| 47 | 75 | Astrocytoma | Female | 25094 | Right | 1.10 | 2.13 |
| 48 | 75 | GBM | Female | 60841 | Left | 0.82 | 1.61 |
| 49 | 18 | Astrocytoma | Male | 5641 | Right | 0.63 | 0.67 |
| 50 | 56 | GBM | Male | 45310 | Left | 1.11 | -1.90 |
| 51 | 28 | Astrocytoma | Male | 62731 | Left | 1.48 | 4.12 |
| 52 | 70 | Astrocytoma | Male | 102302 | Left | 1.01 | 1.51 |
| 53 | 44 | Astrocytoma | Male | 15861 | Right | 0.70 | 4.20 |
| 54 | 66 | GBM | Female | 6054 | Left | 1.96 | -0.93 |
| 55 | 18 | Cortical dysplasia | Female | 21027 | Left | 0.22 | 1.02 |
| 56 | 70 | GBM | Female | 38524 | Right | Could not localize the LMFG ROI | |
| 57 | 67 | GBM | Male | 199912 | Left | 0.97 | 1.62 |

| | | | | | | | |
|---|---|---|---|---|---|---|---|
| 58 | 52 | GBM | Female | 28292 | Left | 1.23 | 2.33 |
| 59 | 34 | Astrocytoma | Female | 278688 | Left | 2.63 | 3.78 |
| 60 | 51 | Gliosis | Male | 15281 | Left | 1.24 | 4.17 |
| 61 | 60 | Oligodendroglioma | Female | 17316 | Left | 0.75 | 3.06 |
| 62 | 25 | Astrocytoma | Male | 74908 | Left | 0.64 | 5.19 |
| 63 | 31 | Oligodendroglioma | Male | 66304 | Left | 0.96 | 3.86 |
| 64 | 59 | Glioma | Male | 77713 | Right | 2.33 | 1.60 |
| 65 | 24 | Oligodendroglioma | Female | 8505 | Left | 2.48 | 0.83 |
| 66 | 59 | Astrocytoma | Female | 12096 | Right | 1.62 | 5.76 |
| 67 | 38 | AVM | Male | 19622 | Left | 1.27 | -0.55 |
| 68 | 59 | GBM | Male | 116479 | Left | 3.04 | 2.33 |
| 69 | 62 | Oligodendroglioma | Female | 25170 | Left | 2.69 | 1.40 |
| 70 | 45 | Astrocytoma | Male | 110326 | Left | 1.38 | 1.80 |
| 71 | 27 | Cavernous malformation | Male | 9046 | Right | 0.29 | 2.43 |
| 72 | 70 | Astrocytoma | Male | 19704 | Left | 1.75 | 2.21 |
| 73 | 47 | Astrocytoma | Female | 85962 | Right | 1.50 | -0.14 |
| 74 | 48 | GBM | Male | 129643 | Left | Could not localize the LMFG ROI | |
| 75 | 21 | AVM | Female | 9459 | Left | 0.93 | 0.60 |
| 76 | 48 | Astrocytoma | Female | 90718 | Left | 2.62 | 3.57 |
| 77 | 26 | Astrocytoma | Male | 12604 | Right | 1.66 | 3.46 |
| 78 | 40 | AVM | Female | 7833 | Left | 1.76 | 5.68 |
| 79 | 36 | Oligodendroglioma | Female | 43092 | Right | 2.00 | 4.08 |
| 80 | 26 | Gliosis | Male | 17726 | Left | 2.65 | 5.99 |
| 81 | 28 | Cortical dysplasia | Female | 39850 | Right | 2.72 | 1.05 |
| 82 | 40 | AVM | Male | 31740 | Right | Could not localize the LMFG ROI | |
| 83 | 53 | Astrocytoma | Male | 48438 | Right | 0.75 | 2.79 |
| 84 | 35 | AVM | Male | 10130 | Left | 1.23 | 2.76 |
| 85 | 40 | Oligodendroglioma | Female | 12465 | Left | 1.74 | 4.95 |
| 86 | 55 | GBM | Female | 129994 | Right | Could not localize the LMFG ROI | |
| 87 | 42 | GBM | Male | 192798 | Left | 0.76 | 0.83 |
| 88 | 76 | GBM | Male | 1538 | Left | Could not localize the LMFG ROI | |
| 89 | 65 | GBM | Female | 102966 | Right | 1.23 | -0.42 |

| | | | | | | | |
|---|---|---|---|---|---|---|---|
| 90 | 28 | Oligodendroglioma | Male | 37022 | Left | 3.13 | 0.33 |
| 91 | 46 | DNET | Female | 9364 | Left | 2.60 | 5.74 |
| 92 | 79 | GBM | Female | 35746 | Left | 0.67 | -1.79 |
| 93 | 76 | GBM | Male | 41761 | Left | 1.63 | 5.10 |
| 94 | 65 | AVM | Male | 33915 | Left | 1.58 | 2.40 |
| 95 | 24 | Glioma | Male | 16251 | Right | 1.55 | 1.89 |
| 96 | 39 | Astrocytoma | Male | 133291 | Right | 0.48 | 2.13 |
| 97 | 41 | Glioma | Male | 33495 | Left | 0.95 | 2.81 |
| 98 | 63 | GBM | Male | 16919 | Left | 0.94 | 1.33 |
| 99 | 53 | Oligodendroglioma | Female | 64556 | Left | 0.66 | 7.54 |
| 100 | 33 | AVM | Female | 15557 | left | 1.19 | 0.33 |
| 101 | 43 | Oligodendroglioma | Female | 23096 | Left | 2.56 | 1.61 |
| 102 | 42 | Astrocytoma | Male | 102665 | Left | 0.69 | 2.98 |
| 103 | 35 | Astrocytoma | Male | 47252 | Left | 1.21 | 2.74 |
| 104 | 40 | Astrocytoma | Female | 26814 | Left | 1.88 | 1.30 |
| 105 | 62 | GBM | Male | 83984 | Left | 0.55 | 5.54 |
| 106 | 75 | GBM | Male | 96475 | Left | Could not localize the LMFG ROI | |
| 107 | 34 | Oligodendroglioma | Male | 71566 | Left | 2.24 | 5.57 |
| 108 | 22 | Astrocytoma | Male | 124642 | Right | 0.96 | 0.14 |
| 109 | 44 | PLNTY tumor | Male | 14179 | Left | 1.54 | -1.59 |
| 110 | 49 | Glioma | Female | 23800 | Left | 1.30 | 1.44 |
| 111 | 23 | Glioneuronal tumor | Male | 3871 | Right | 1.65 | 1.95 |
| 112 | 55 | GBM | Female | 85363 | Right | 1.11 | 3.08 |
| 113 | 23 | Astrocytoma | Male | 30201 | Left | 2.25 | 0.35 |
| 114 | 30 | Oligodendroglioma | Male | 220690 | Left | Data discarded due to technical issue | |

**B.** Neurotypical participants

| Cumulative Participant ID | Age | Gender | Cross-validated T>AFP *t*-value | Cross-validated P>AFT *t*-value |
|---|---|---|---|---|
| 1 | 21 | Female | 0.36 | 4.36 |
| 2 | 19 | Male | 0.53 | 3.28 |
| 3 | 22 | Male | 0.37 | -2.13 |

| | | | | |
|---|---|---|---|---|
| 4 | 22 | Male | 2.01 | 1.57 |
| 5 | 20 | Male | 1.13 | 0.32 |
| 6 | 19 | Male | 0.66 | -0.41 |
| 7 | 18 | Male | 0.76 | 2.24 |
| 8 | 24 | Female | 2.48 | 2.43 |
| 9 | 26 | Male | 2.48 | 1.62 |
| 10 | 20 | Female | 1.16 | 1.53 |
| 11 | 20 | Female | 1.55 | 5.17 |
| 12 | 19 | Female | 2.28 | 0.64 |
| 13 | 21 | Female | 1.17 | 1.29 |
| 14 | 19 | Female | 1.56 | -0.47 |
| 15 | 23 | Male | 2.04 | -1.04 |
| 16 | 20 | Female | 1.80 | 3.46 |
| 17 | 21 | Female | 0.99 | 0.08 |
| 18 | 19 | Female | 1.29 | 3.09 |
| 19 | 20 | Male | 0.52 | 4.92 |
| 20 | 22 | Female | 1.03 | 5.11 |
| 21 | 19 | Female | 0.66 | 1.68 |
| 22 | 19 | Female | 1.07 | 2.19 |
| 23 | 21 | Female | 2.06 | -0.14 |
| 24 | 21 | Female | 1.20 | 4.84 |
| 25 | 21 | Male | 0.64 | 6.07 |
| 26 | 19 | Male | 1.33 | 4.94 |
| 27 | 19 | Female | 0.36 | 4.19 |
| 28 | 20 | Female | 1.23 | 2.33 |
| 29 | 20 | Male | Could not localize the LMFG ROI | |

| | | | | |
|---|---|---|---|---|
| 30 | 21 | Male | 0.43 | 2.29 |
| 31 | 23 | Male | 1.94 | 2.79 |
| 32 | 19 | Female | 1.37 | 2.51 |
| 33 | 21 | Male | 1.73 | -0.11 |
| 34 | 21 | Female | 1.57 | 3.56 |
| 35 | 27 | Female | 1.42 | 3.95 |
| 36 | 20 | Male | 2.53 | -4.85 |
| 37 | 26 | Male | Could not localize the LMFG ROI | |
| 38 | 57 | Female | 0.96 | -0.26 |
| 39 | 43 | Female | 1.26 | 2.99 |
| 40 | 20 | Male | Could not localize the LMFG ROI | |
| 41 | 20 | Female | 2.20 | 5.42 |
| 42 | 24 | Male | 0.67 | 0.79 |
| 43 | 30 | Male | 0.58 | 7.56 |
| 44 | 23 | Female | 1.27 | 3.73 |
| 45 | 21 | Female | 0.94 | 5.28 |
| 46 | 23 | Female | 1.04 | -1.75 |
| 47 | 24 | Male | 0.83 | 2.36 |
| 48 | 20 | Female | 0.93 | 0.49 |
| 49 | 21 | Male | 1.55 | 1.17 |
| 50 | 20 | Male | 0.56 | 3.28 |
| 51 | 22 | Male | 0.93 | 3.2 |
| 52 | 21 | Female | 0.48 | 5.47 |
| 53 | 22 | Male | 1.53 | 3.83 |
| 54 | 21 | Female | 1.53 | -2.67 |
| 55 | 21 | Male | 1.75 | 7.64 |
